# Translation stalling proline motifs are enriched in slow-growing, thermophilic, and multicellular bacteria

**DOI:** 10.1101/2021.10.05.463052

**Authors:** Tess E. Brewer, Andreas Wagner

## Abstract

Rapid bacterial growth depends on the speed at which ribosomes can translate mRNA into proteins. mRNAs that encode successive stretches of proline can cause ribosomes to stall, substantially reducing translation speed. Such stalling is especially detrimental for species that must grow and divide rapidly. Here we focus on di-prolyl motifs (XXPPX) and ask whether their incidence varies with growth rate. To find out we conducted a broad survey of such motifs in >3000 bacterial genomes across 36 phyla. Indeed, fast-growing species encode fewer motifs than slow-growing species, especially in highly expressed proteins. We also found many di-prolyl motifs within thermophiles, where prolines can help maintain proteome stability. Moreover, bacteria with complex, multicellular lifecycles also encode many di-prolyl motifs. This is especially evident in the slow-growing phylum Myxococcota. Bacteria in this phylum encode many serine-threonine kinases, and many di-prolyl motifs at potential phosphorylation sites within these kinases. Serine-threonine kinases are involved in cell signaling and help regulate developmental processes linked to multicellularity in the Myxococcota. Altogether, our observations suggest that weakened selection on translational rate, whether due to slow or thermophilic growth, may allow di-prolyl motifs to take on new roles in biological processes that are unrelated to translational rate.

## Introduction

Translation is a fundamental process common to all known forms of life. Cells invest huge amounts of resources into translation. For example, in fast-growing bacterial species like *E. coli* protein synthesis can account for over 50% of a cell’s total energy budget [1]. What is more, rapid bacterial growth depends on the speed of translation. Specifically, fast-growing bacteria maintain high concentrations of ribosomes, and these ribosomes elongate proteins rapidly during protein synthesis [2]. In addition, bacterial genome characteristics that are correlated with growth rate (rRNA and tRNA gene copy number, as well as codon usage bias) all influence the rate of translation [3–6].

Multiple factors can negatively impact the translation rate and cause ribosomes to pause or ‘stall’ during elongation. These factors include the presence of uncharged tRNAs, rare codons in a translated mRNA, and even specific amino acids encoded by mRNA [7]. Among these amino acids proline stands out. Proline is slow to form peptide bonds due to its structural rigidity and unique status as an N-alkylamino acid [8, 9]. This structural rigidity can contribute to the formation of special secondary structures, like the poly-proline II helix [10, 11], which is associated with the binding domains of signaling proteins [12, 13]. Successive stretches of prolines also cause ribosomes to pause translation. The length of this pause—the ‘strength’ of the ribosome ‘stall’—depends on the amino acids surrounding the proline stretch [14], the location of the sequence causing the stall within a protein [15], and the translation initiation rate [16].

A special translation factor exists to resolve proline-induced ribosomal stalls. In bacteria this protein is called translation elongation factor P (EFP). EFP is a tRNA mimic that binds to the ribosome between the peptidyl and exit sites [17]. When bound to the ribosome, EFP uses a conserved amino acid residue to interact with the peptidyl-transferase center and to accelerate the formation of proline-proline peptide bonds [17]. In many species, this conserved amino acid must be post-translationally modified for EFP to efficiently alleviate stalling [18–20]. The importance of EFP and its mitigation of ribosome stalling is underscored by the strong phenotypes caused by its loss. These include diminished growth rate [20–24], loss of motility [25], loss of virulence[20, 22, 24], reduced antibiotic resistance [24, 26], and in some cases, cell death [27].

While EFP can reduce the impact of proline-induced ribosomal stalling, EFP cannot completely eliminate these stalls. Ribosomal profiling shows that *E. coli* ribosomes still pause at proline residues, albeit much more briefly than in EFP knockout mutants [15]. Indeed, recent work has directly shown that proline motifs lead to ribosomal pausing in wild-type *E. coli* [28]. Protein evolution may have exploited such unavoidable stalling. For example, in *E. coli* di-prolyl motifs often occur at the beginning of complex protein domains, and may provide additional time for translational regulation, protein folding, or membrane insertion [29]. Indeed, *E. coli* appears to prefer rare proline codons for such motifs, effectively lengthening the stall phenotype in these regions [28].

Because EFP cannot fully alleviate proline-induced stalling [15, 28], one would expect that stalling motifs are subject to natural selection, and especially so in fast-growing species under high pressure to maximize their translation rate [2, 4]. Indeed, di-prolyl sequences occur less frequently than expected by chance in the fast-growing *E. coli*, where highly expressed proteins are especially depleted in these motifs [29]. EFP itself is optimized for high expression in fast-growing bacteria [30], reflecting its importance in maintaining high growth rates.

Slow-growing bacteria are under reduced selection for translational speed [31]. Their genomes have reduced codon usage bias and encode fewer tRNA and rRNA gene copies than their fast-growing counterparts [5, 32]. Therefore, we wondered whether the di-prolyl motifs that can cause ribosome-stalling would be more widespread in slow-growing bacteria. The prevalence of such motifs is unknown outside few well-studied bacterial species, including *E. coli* [29], *S. enterica* [33], *Bacillus subtilis* [23], and several Actinobacteria species [34].

We quantified the occurrence of di-prolyl motifs across more than 3000 bacterial genomes from 35 phyla and found that these motifs are more abundant in genomes with high GC content. This is not surprising, because proline codons are cytosine rich. More importantly, we found that these motifs were more abundant in species with slow predicted growth rates when we controlled for GC content. Di-prolyl motifs are also more abundant in thermophiles, and in species with complex life cycles that involve a multicellular life stage. They are especially abundant in the serine-threonine protein kinases of multicellular species, which are involved in signaling and developmental programs.

## Materials & Methods

### Analysis of Bacterial Genomes

We downloaded 3265 bacterial genomes from the Integrated Microbial Genomes (IMG) database[35], selecting only one genome per Average Nucleotide Identity (ANI) cluster to reduce bias towards highly studied species. This procedure yielded approximately one representative genome from each species in the database, although we included multiple genomes from a species if the ANI between genomes was below the typical cutoff for species level (less than 96.5, 11 species). We used CheckM [36] to evaluate the quality of these genomes, only retaining those which were at least 90% complete and contained less than 5% contamination, based on their distribution of highly conserved single copy genes. We also re-assigned taxonomy to the whole dataset using the Genome Taxonomy Database and GTDB-Tool kit (GTDB-Tk) version 0.2.2 [37], and removed any genomes that could not be assigned to a phylum (three genomes). We counted the occurrence of di-prolyl motifs (XXPPX, where X designates any amino acid) in every protein encoded in each genome, using custom python scripts. We count polyproline motifs (XPPPX) as multiple di-prolyl motifs. For example, AAPPPA would be represented as the two di-prolyl motifs AAPPP and APPPA. We followed this procedure because from the point of view of the ribosome such motifs represent independent proline-proline bond formation reactions. The identity of the amino acids surrounding successive prolines (Xs) has been shown to impact the severity or length of the resulting ribosomal stall [14, 29]. We classified each di-prolyl motif according to its predicted stall severity, from weak to medium to strong, using a key derived from a mixture of *in vitro* and *in vivo* data [29].

We verified the presence of at least one EFP homolog in nearly every genome using the `hmmscan` function of HMMER version 3.3.2[38] to search for Pfam PF01132. Only the genome of *Aquaspirillum serpens* did not encode a known EFP homolog. However, because this genome is not fully complete (estimated completeness 98.27% by CheckM), and because it encodes the EFP modification protein earP [20], it likely does encode EFP.

Next, we estimated the doubling time associated with each genome using the codon usage bias (CUB) based R package gRodon version 1.8.0 [32]. This package calculates estimated doubling times by comparing the CUB from a set of genes expected to be highly expressed in fast-growing species (we used ribosomal proteins) to the background codon usage of the genome, with the expectation that fast-growing species use codons corresponding to the most abundant tRNAs for maximum translational rate in growth related genes. This metric provides a good approximation for a species’ doubling time in both whole genomes and metagenomic samples [32].

For a subset of our genomes, we retrieved experimentally measured doubling times from the literature (see **Supplementary Dataset S1** for all corresponding citations), with a large proportion of this data coming from a recently compiled database on bacterial phenotypes [39]. Because we matched genomes with phenotypic data by species name, the genomes in our dataset are not necessarily from the exact strain for which doubling times were measured, but they will generally be closely related. Wherever more than one entry for the same species existed in the phenotypic database, we used the fastest recorded doubling time for our analyses. Wherever our genome data set harbored multiple genomes for the same species, we used the genome with the fastest CUB-predicted doubling time. We found good agreement between doubling times predicted by gRodon and measured doubling times, especially when only mesophilic species were considered (species with measured doubling times: n = 301, Pearson‘s rho = 0.33, p <0.0001; mesophiles only: n = 202, Pearson’s rho = 0.44, p < 0.0001, **Figure S1**). In addition, regardless of how accurately CUB reflects measured doubling times, it is valuable to analyze CUB in this context, because it reflects a species’ investment in optimizing its translation rate.

In order to calculate the median expected expression level of genes, we first used ENCprime [40] to calculate the CUB for each individual gene (represented as KEGG KOs) within our genome dataset. We then ranked each gene in each genome, assigning the highest rank of one to the gene with the highest CUB. We took the median codon bias rank for each gene across all genomes encoding said gene and used these values to approximate the median expression level of each gene across all species in our data set. We used the median codon bias rank to prevent genomes belonging to fast-growing species from overly biasing the results. Based on this calculation, the five genes with the highest predicted expression level encoded elongation factor Tu, chaperonin GroEL, large subunit ribosomal protein L7/L12, small subunit ribosomal protein S1, and elongation factor Ts. These results are consistent with the expectation that genes related to translation and cell growth should be highly expressed across most genomes. All KEGG annotations were provided by IMG using their annotation pipelines [35].

We identified thermophiles and psychrophiles in our dataset by using either 1) IMG-provided temperature ranges, 2) IMG-provided habitat data (e.g., species from hot springs and hydrothermal vents were categorized as thermophiles), or 3) membership in phylogenetic groups with conserved temperature ranges (such as known thermophilic orders like the Thermales or Aquificales). To identify bacteria with a complex, multicellular lifestyle, we consulted a review on multicellularity [41] and reviews on specific bacterial phyla [42–44]. We identified serine-threonine kinases by extracting all proteins which fell within KEGG orthology group K08884.

To identify intrinsically disordered regions (IDRs) within proteins of interest, we used IUPred2A [45] with the ‘long’ option. IUPred predicts IDRs by using statistical potentials to estimate the total pairwise interaction energy between a stretch of consecutive amino acids (30 amino acids in the ‘long’ option). IDRs occur when amino acids cannot form favorable interactions because of low pairwise interaction energy. IUPred computes a ‘disorder score’, and when this score exceeds a value of 0.5 in a protein region, the region is predicted to be disordered. We calculated an average disorder score for each di-prolyl motif by averaging scores across all five amino acids in each motif (IUPred computes a disorder score for each individual amino acid). **Supplementary Datasets S1** and **S2** contain information on all genomes and all proteins we analyzed, respectively.

### Statistical methods

One potentially confounding factor in our analysis is that proline codons are cytosine rich (CCU, CCC, CCA, and CCG), which implies that di-prolyl motifs are inherently more likely to occur in genomes with high GC content. Indeed, the number of coding GC base pairs and the number of di-prolyl motifs are very strongly correlated for genomes in our dataset (Pearson’s rho = 0.94, p < 0.0001). Because of this correlation, when examining individual proteins and their di-prolyl content, we controlled for the GC content of their encoding gene. We also controlled for protein length, because long proteins may contain more di-prolyl motifs than short proteins by chance alone. We controlled for both quantities by dividing the total number of nucleotides encoding the di-prolyl motifs (each motif is five amino acids long and thus encoded by 15 base pairs) by the ratio of the total number of base pairs in the gene to the number of GC base pairs in the gene. That is, we preformed all analyses of di-prolyl motifs within genes with the quantity

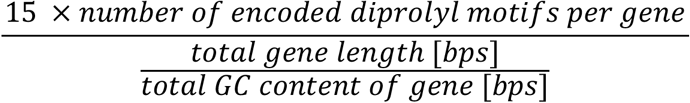

Another potential confounding factor in our analyses is that GC content and other genomic characteristics are correlated across bacterial phylogenies. In other words, closely related bacteria are more likely to have similar GC content—and thus di-prolyl content—than those that are distantly related. To account for such phylogenetic dependence, we created a phylogenetic tree using 43 concatenated conserved marker genes generated by CheckM [36]. We aligned these sequences using MUSCLE version 3.8.31 [46], and built the phylogenetic tree with FastTree version 2.1.10 [47], using the archaeon *Haloquadratum walsbyi* as an outgroup (NCBI accession number: GCA_000009185). We used this tree for all subsequent phylogeny-dependent statistical methods.

Next, we calculated Pagel’s λ [48] using the phylosig function from the `phytools` R package, version 0.6.99 [49]. Pagel’s λ is a measure of the phylogenetic dependence of a trait. A value of λ = 0 indicates that the trait evolved independently of phylogeny, while λ = 1 indicates strong phylogenetic dependency. This calculation confirmed that GC content shows strong phylogenetic dependency (λ = 0.99, p < 0.0001). Therefore, we controlled for phylogeny in all our genome-based analyses.

To this end, we used phylogenetic generalized least squares (PGLS) to measure the contribution of individual genomic characteristics to the prevalence of di-prolyl motifs within our genomes. We also used a phylogenetic ANOVA to analyze differences between groups in our dataset. For the PGLS, we used the `pgls` function in the R package caper, version 1.0.1[50] and for the phylogenetic ANOVA we used the `phylANOVA` function from the R package phytools version 0.7-70 [49]. We performed all statistical analyses and plotting in R version 3.6.2 and created all plots using ggplot2 version 3.3.3 [51].

## Results

### Species with large, high GC genomes have many di-prolyl motifs

We quantified the frequency of di-prolyl motifs (XXPPX, where X designates any amino acid) in all proteins within a set of >3000 bacterial genomes from 31 different phyla. While all di-prolyl motifs (PP) cause ribosomal stalling, the surrounding amino acids (X) can influence the severity of the stall [14]. To assign a stall ‘strength’ to each di-prolyl motif, ranging from strong to medium to weak, we used a published key that has been compiled from *in vivo* and *in vitro* proteomic experiments [29].

We found that the number of di-prolyl motifs in each genome varied broadly throughout our dataset. They range from a maximum of 17,841 (1.95 motifs per protein) in *Nannocystis exedens* (a Myxobacterium with a complex lifecycle) to a minimum of 86 (0.15 motifs per protein) in *Mycoplasma cloacale*, a poultry-associated pathogen from the family *Mycoplasmataceae*. The genome of *N. exedens* also contained the most ‘strong’ di-prolyl motifs at 9665 (1.05 strong motifs per protein), while *Mesoplasma coleopterae*, another pathogen from the *Mycoplasmataceae*, had the fewest strong motifs at just 17 (0.02 strong motifs per protein). In general, we found that phyla with large, high GC genomes had the highest number of motifs (Actinobacteria, Planctomycetota, and Myxococcota), while those with small, low GC genomes had the fewest motifs (Fusobacteria, Campylobacterota, and Thermotogota, **Figure 1**). This is not surprising because proline codons are cytosine rich (CCU, CCC, CCA, and CCG), making di-prolyl motifs inherently more likely to occur in large genomes with high GC content. Next, we asked whether any genome-derived characteristics besides GC content and genome size influence the frequency of di-prolyl motifs. When quantifying the influence of these characteristics on the prevalence of di-prolyl motifs, we must control for the shared evolutionary history of our study taxa. For example, closely related genomes are much more likely to have similar GC content, and therefore similar numbers of di-prolyl motifs, than expected by chance (Pagel‘s λ = 0.99, see Methods). In addition, we needed a method that could account for the highly correlated relationship between the frequency of di-prolyl motifs and genomic GC content. To disentangle the contributions of these and other characteristics, while also controlling for phylogeny, we used a phylogenetic generalized linear model (PGLS). This statistical method uses a phylogenetic tree to control for phylogenetic relatedness, essentially ‘down-weighting’ similar observations that originate from closely related species [52], while also accounting for co-correlated variables.

**Figure 1:**
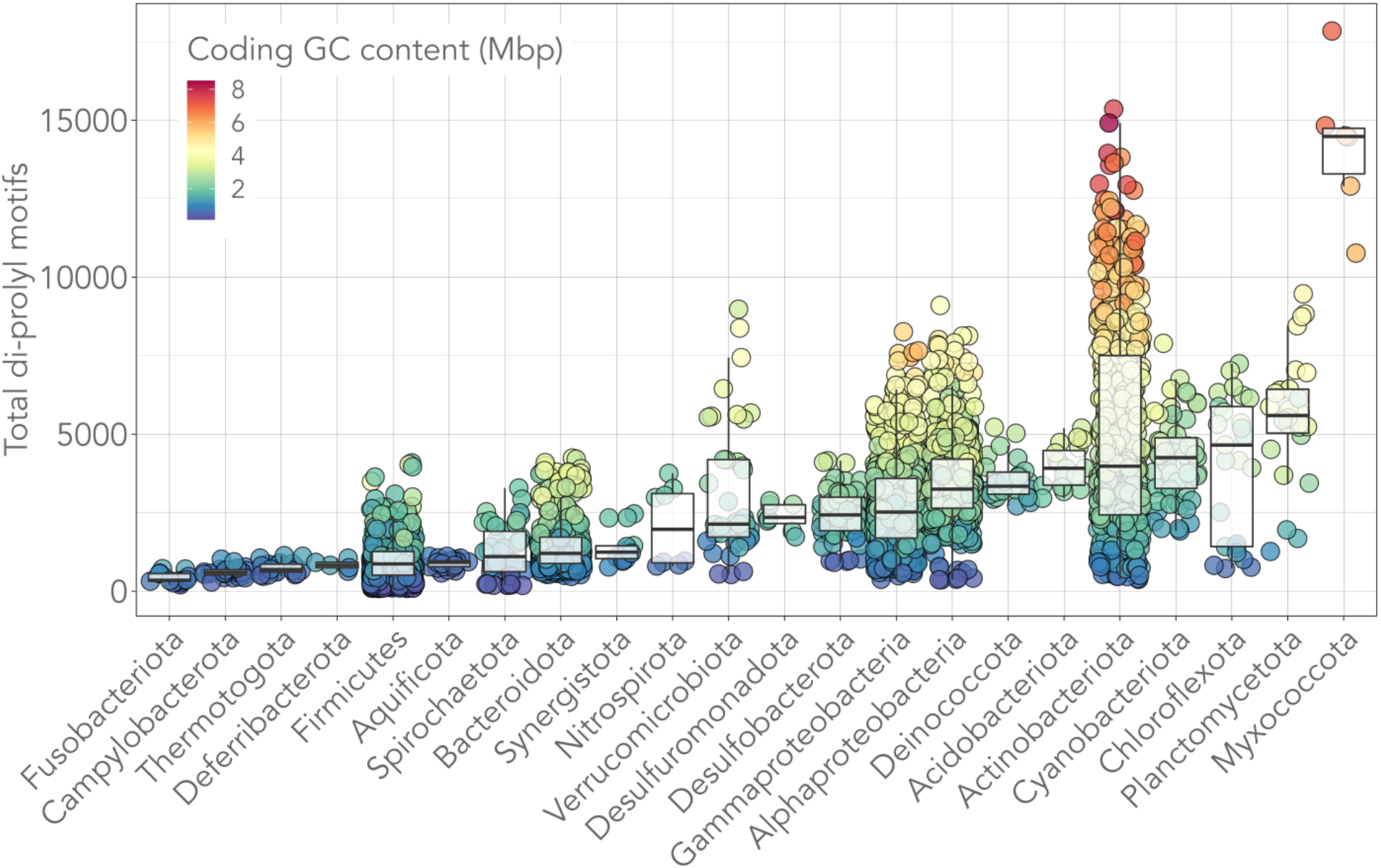
Di-prolyl motifs occur frequently in bacteria with large, high GC genomes. Phyla whose genomes have a low average GC coding content (Fusobacteriota 0.8 Mbp, Campylobacterota 0.7 Mbp, Thermotogota 0.8 Mbp) encode fewer di-prolyl motifs than phyla with high average GC coding content (Actinobacteria 3.2 Mbp, Planctomycetota 3.2 Mbp, Myxococcota 6.3 Mbp). Only phyla represented by at least five genomes in our dataset are shown. Each circle represents one genome. Taxonomy was assigned using GTDB (Genome Database Taxonomy; see Methods). The phylum Proteobacteria was broken down into its corresponding classes and all Firmicutes-adjacent phyla were combined.

### Thermophiles and microbes with complex life cycles have high levels of di-prolyl motifs

The structural rigidity of proline does not just lead to slow peptide bond formation. It can also reduce the conformational freedom of polypeptide chains, leading to increased thermo-stabilization [53]. Additionally, like slow-growing species, thermophiles are thought to experience weaker selection on growth-associated-traits than mesophiles. This is because high temperatures cause higher rates of catalysis and tRNA diffusion [5]. In other words, compared to mesophiles growing at the same speed, thermophiles need to invest less in optimizing growth-associated traits like rRNA and tRNA gene copies and codon usage bias. For these reasons, we hypothesized that di-prolyl motifs may be more abundant in thermophilic bacteria. Testing this hypothesis is complicated by the fact that thermophiles have smaller genomes and shorter proteins than mesophiles [54]. With this in mind, we first performed a phylogenetic ANOVA which confirmed that thermophiles encode more di-prolyl motifs per Mbp of GC coding content than mesophiles (thermophilic mean = 1389 di-prolyl motifs per coding GC Mbp, mesophilic mean = 1150 di-prolyl motifs per coding GC Mbp; Phylogenetic ANOVA, p-value < 0.05, **Figure 2A**). Next, we performed a PGLS to verify that thermophiles encoded more di-prolyl motifs when total GC coding sequence size was controlled for. We again found that thermophiles encode significantly more di-prolyl motifs than mesophiles (**PGLS A**, p < 0.0001, b_*thermophiles*_ = 0.023, b = regression slope, **Supplemental Table 1**). Psychrophiles did not differ from mesophiles in this respect (**PGLS A**, p = 0.53), although this could be due to their comparatively poor representation in our dataset (mesophiles n = 2892, thermophiles n = 304, psychrophiles n = 56).

**Figure 2:**
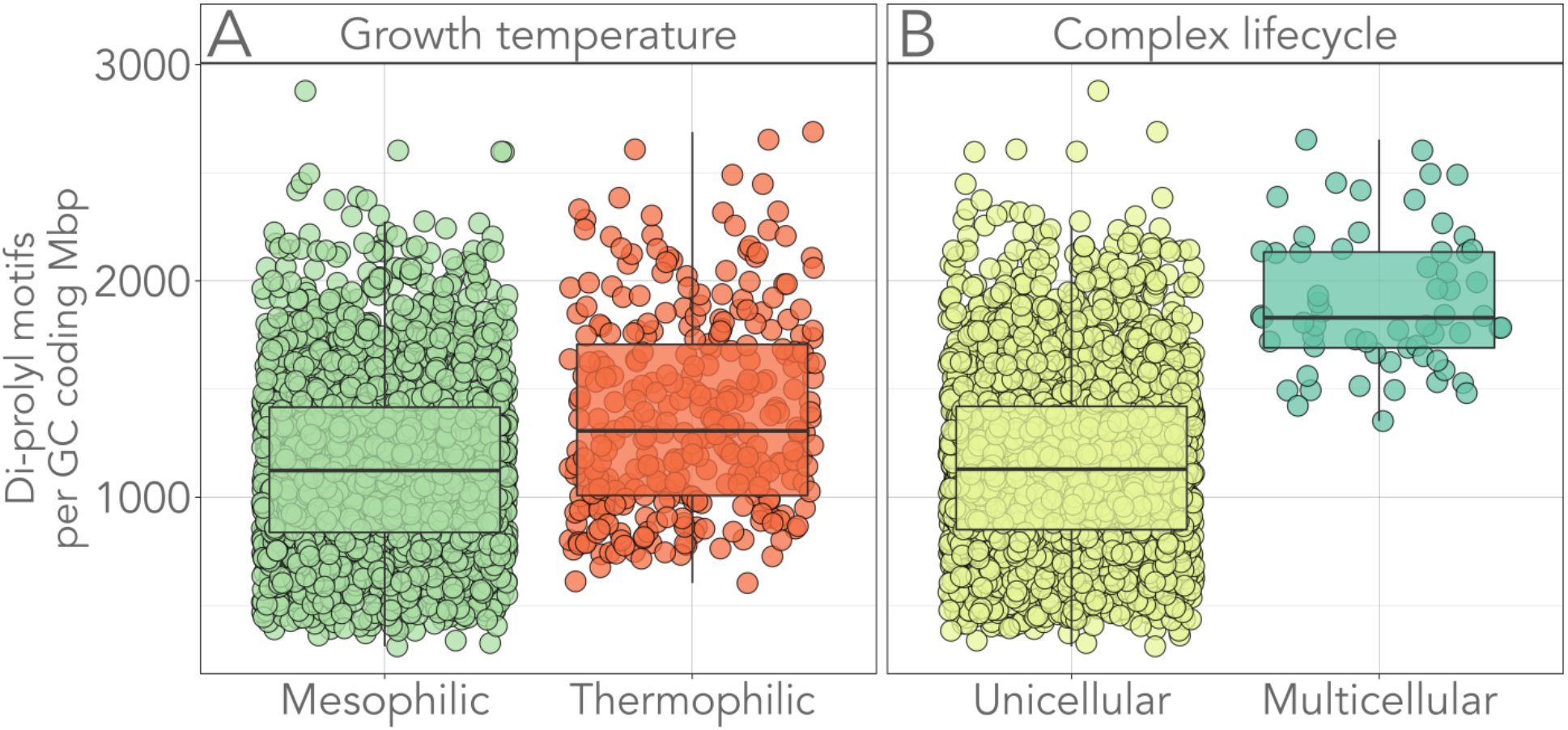
Thermophilic and multicellular bacteria encode many di-prolyl motifs per GC coding Mbp. **A)** Thermophile genomes encode significantly more di-prolyl motifs per GC coding Mbp than mesophiles (Phylogenetic ANOVA, p-value < 0.05). Thermophiles are represented by 304 genomes, mesophiles by 2892 genomes: thermophilic mean = 1389 di-prolyl motifs per GC coding Mbp, mesophilic mean = 1150 di-prolyl motifs per GC coding Mbp. **B)** The genomes of species with a complex, multicellular lifecycle encode significantly more di-prolyl motifs per GC coding Mbp than their unicellular counterparts (Phylogenetic ANOVA, p-value ≤ 0.001). Multicellular and unicellular species are represented by 61 and 3191 genomes, respectively: multicellular mean = 1902 motifs per GC coding Mbp, unicellular mean = 1157 motifs per GC coding Mbp.

Anecdotal evidence from our initial data exploration showed that the Myxococcota, which are well-known for their ability to form multicellular fruiting bodies, have the most di-prolyl motifs among all taxonomic groups we examined (**Figure 1**), despite not having the highest overall GC content. Myxococcota and other bacteria with complex life cycles rely on cell-cell signaling to orchestrate their developmental programs [55]. Proline-rich regions often occur in the binding domains of signaling proteins, where they mediate protein-protein binding in a highly specific yet reversible manner [13]. Therefore, we wondered whether the genomes of bacteria with complex lifecycles are generally more likely to harbor many di-prolyl motifs.

To find out, we performed a phylogenetic ANOVA to find out whether bacteria known for multicellular behavior encoded more di-prolyl motifs per GC coding Mbp. This is indeed the case (Phylogenetic ANOVA, p ≤ 0.001, multicellular mean = 1902 di-prolyl motifs per coding GC Mbp, unicellular mean = 1157 di-prolyl motifs per coding GC Mbp, **Figure 2B**). Next, we modified our PGLS model to include multicellularity as an added variable. In addition, we also included temperature class in the model, because some multicellular bacteria are thermophilic, for example the filamentous thermophile *Ardenticatena maritima*. We again found that multicellular bacteria have significantly more di-prolyl motifs than unicellular bacteria, after controlling for total GC coding content and growth temperature class (**PGLS B**, p < 0.01, b_unicellular_ = −0.016). We also asked whether the elevated di-prolyl content of motifs in multicellular bacteria was solely attributable to the Myxococcota, because this phylum contain the most di-prolyl motifs of any taxonomic group (**Figure 1**). This was not the case: when we removed Myxococcota genomes from consideration, multicellular genomes still encoded more di-proyl motifs than unicellular genomes (**PGLS C**, p < 0.05, b_unicellular_ = −0.015).

### Slow-growing species encode more di-prolyl motifs than fast-growing species

Because di-prolyl motifs can negatively impact translation rate [15, 28], we wanted to find out whether selection for translational speed would impact the number of di-prolyl motifs in a genome. Ideally, we would answer this question using experimentally measured growth rates. Unfortunately, this information is not widely available—we could find experimentally validated growth rates for only 301 species in our dataset. As an alternative, we calculated the predicted growth rate of each species in our dataset using a codon usage bias (CUB) centered method, gRodon [32]. CUB refers to the tendency of species to use codons that correspond to the most abundant tRNAs in highly expressed genes. In doing so, these species can increase their translational rate by accelerating tRNA turnover at the ribosome [5]. The degree of CUB in genes encoding ribosomal proteins is well correlated with experimentally measured growth rates in mesophilic species [5, 32, 56] (see Methods, **Figure S1)**. In addition, regardless of how strongly CUB-predicted growth rates and experimentally measured growth rates may be correlated, using CUB itself in this analysis is valuable because CUB represents the degree of investment a species has made towards maximizing translational rate [5].

Along with predicted growth rates, we also included two other growth-associated traits in this PGLS model. These are the copy numbers of tRNA and rRNA genes. One strategy that fast-growing bacteria use to translate proteins rapidly is to ensure that their pool of charged tRNA does not become limiting. Fast-growing species thus often encode multiple copies of the most common tRNA genes [57]. Similarly, fast-growing species tend to encode multiple rRNA gene copies to boost the rate at which rRNA molecules—and consequently ribosomes—are synthesized [58]. When included in our PGLS model, all three growth-associated traits had a significant impact on the number of di-prolyl motifs in a genome, although the significance of rRNA gene copies was weak (**PGLS D**; predicted doubling time p < 0.0001 b = 0.016, tRNA gene copies p < 0.005 b = −0.030, rRNA gene copies p < 0.1, b = −0.003). The direction of the impact of these traits was uniform. That is, characteristics linked to slow-growth (slower predicted doubling times, fewer tRNA gene copies, and fewer rRNA gene copies) were all associated with more di-proly motifs (model details in **Supplemental Table 1**).

Using the growth-associated traits of a representative slow (*Methylomagnum ishizawai*; predicted doubling time = 124 hrs, tRNA gene copies = 51, rRNA gene copies = 2) and fast growing species (*Propionigenium maris*; predicted doubling time = 0.11 hrs, tRNA gene copies = 103, rRNA gene copies = 5) in the equation supplied by the PGLS D model, we found that slow-growth traits resulted in a 14% increase in the number of di-prolyl motifs within a genome, irrespective of GC content (**Figure 3**). This can yield a substantial total increase at a high GC content. For example, at a protein-coding GC content of 8 Mbp, slow-growth associated traits yielded an additional 1283 di-prolyl motifs (**Figure 3C**). The effect of slow-growth associated traits on di-prolyl motifs is independent of growth temperature and multicellularity (**Supplemental Table 1**).

**Figure 3:**
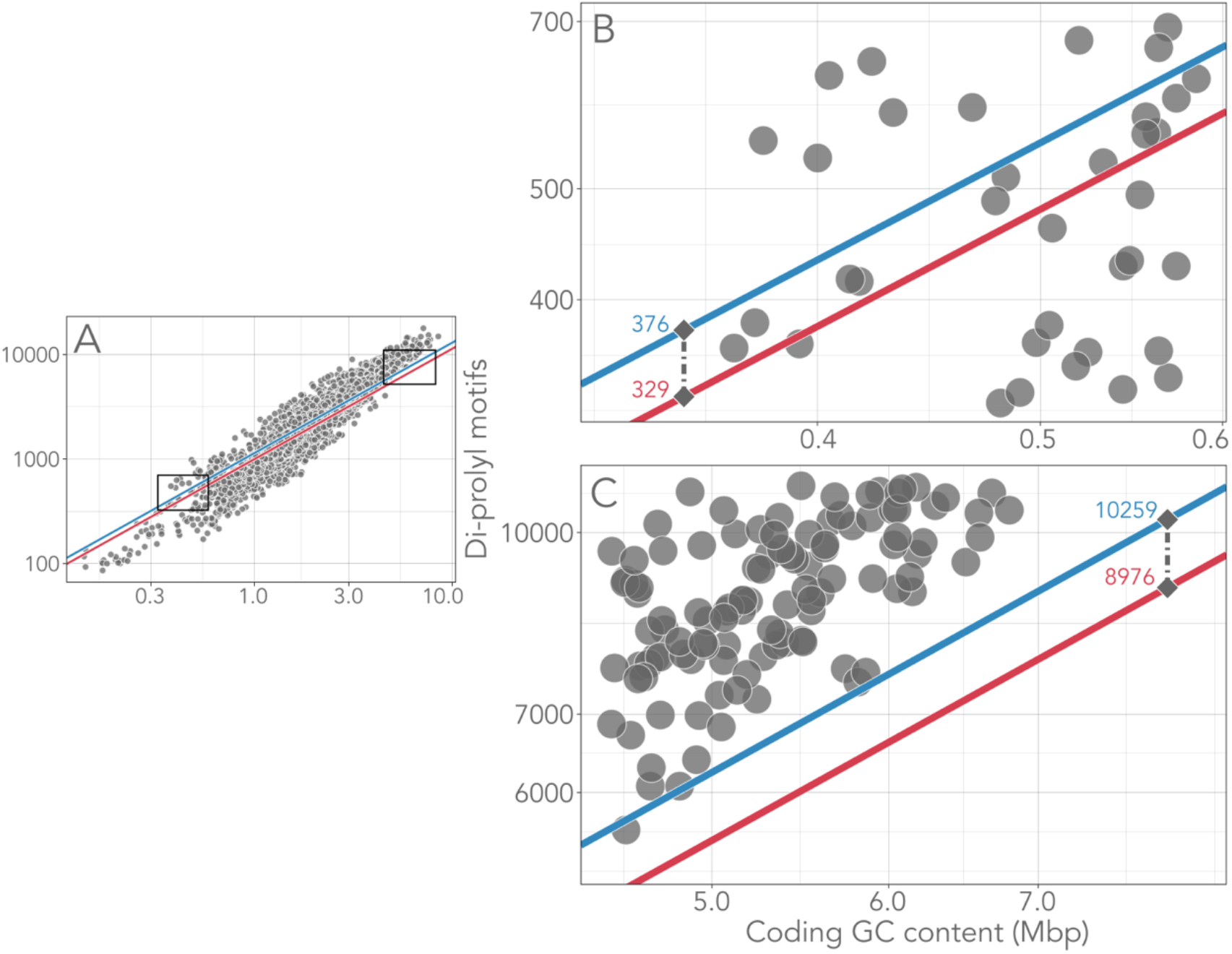
Predicted doubling time and other growth-associated traits significantly affect the abundance of di-prolyl motifs encoded by genomes. **A:** While GC content is the primary determinant of the number of di-prolyl motifs in a genome (**PGLS D** p < 0.0001, regression slope b =1.056), predicted doubling time, tRNA gene copies, and rRNA gene copies also have a significant impact (**PGLS D**; predicted doubling time p < 0.0001 b = 0.016, tRNA gene copies p < 0.005 b = −0.030, rRNA gene copies p < 0.1, b = −0.003). The two diagonal lines represent the number of PGLS-predicted di-prolyl motifs, as calculated from the growth-associated traits of two representative organisms in our dataset. Specifically, the upper blue line represents a prediction based on one of the slowest growing species (*Methylomagnum ishizawai*; predicted doubling time = 124 hrs, tRNA gene copies = 51, rRNA gene copies = 2) while the lower red line represents a prediction based on one of the fastest growing species (*Propionigenium maris*; predicted doubling time = 0.11 hrs, tRNA gene copies = 103, rRNA gene copies = 5). The slow-growth related traits of *Methylomagnum ishizawai* result in a 14% predicted increase in di-prolyl motifs. At low GC content (**B: enlarged view of the lower left box in A**), the total impact of growth-associated traits is low (calculated net increase of only 47 di-prolyl motifs per genome at 0.35 Mbp GC content), while at high GC content (**C: enlarged view of the upper right box in A**) the total net increase is substantial (calculated net increase of 1283 di-prolyl motifs at 8 Mbp GC content). The horizontal and vertical axes are plotted on a logarithmic scale.

While CUB based growth rate metrics predict experimentally measured doubling times well [5, 32] (**Figure S1)**, we wondered whether the statistical associations we had detected would persist if we used experimentally measured doubling times in place of predicted doubling times. We found such data for 301 (9.2%) species in our dataset (see **Supplemental Dataset 1** for details). When we repeated our PGLS analysis on this reduced dataset, we found that species with faster experimentally measured growth rates still encoded fewer di-prolyl motifs than slow-growing species, although rRNA gene copies were no longer a significant predictor (**PGLS E**, measured doubling times p < 0.05 b = 0.015, tRNA gene copies p ≤ 0.005 b = −0.129, rRNA gene copies p = 0.440 b = 0.008). Removing thermophiles from this analysis significantly improved this relationship (**PGLS F**, measured doubling times p ≤ 0.001 b = 0.034, tRNA gene copies p < 0.05 b = −0.108, rRNA gene copies p = 0.315 b = 0.011). This may be explained by the increase in growth rate that thermophiles achieve purely from high optimal growth temperatures: unlike mesophiles, rapid thermophilic growth rates do not necessarily reflect enhanced investment in maximizing translational speed [5]. Using the equation supplied by **PGLS F**, the slow-growth associated traits of *Methylomagnum ishizawai* (experimentally measured doubling time = 24 hrs) yielded a 25% increase in di-prolyl motifs over the fast-growing *Propionigenium maris* (experimentally measured doubling time = 0.3 hrs) (**Figure S2).** In sum, both CUB-predicted and experimentally measured growth rates support the notion that fast-growing species encode fewer di-prolyl motifs than slow-growing species.

### Proteins optimized for translational speed contain few di-prolyl motifs, especially in fast-growing species

Our analysis thus far focused on the incidence of di-prolyl motifs in entire genomes, but this incidence may also vary among proteins within a genome. For example, in *E. coli*, highly expressed proteins have fewer di-prolyl motifs than lowly expressed proteins [29]. We wondered whether this link between expression level and di-prolyl motifs exists more generally in the >3000 bacterial genomes we analyzed. To address this question, we used the KEGG (Kyoto Encyclopedia of Genes and Genomes) Orthology database [59], which assigns each gene in our dataset to a KEGG Orthology (KO) group. This classification provides single-source functional annotations which place each gene in a hierarchical classification scheme that ranges from coarse-grained, e.g., ‘genetic information processing‘, to fine-grained, e.g., ‘ribosomal protein-coding’.

Because gene expression data does not exist for the vast majority of our genomes, we used the codon usage bias of individual genes as a proxy for their expression level. Specifically, we ranked each gene based on its overall CUB, where the highest rank of one corresponds to the gene with the strongest bias and highest predicted expression in each genome. We then took the median of this rank for each gene across all genomes. We controlled for the GC content and length of each protein-coding gene in this analysis (see Methods), and only included genes that were present in at least 25% of genomes to reduce bias towards rare genes.

We found that highly expressed proteins (low median CUB rank) contained significantly fewer di-prolyl motifs, an observation that holds both for all motifs (Spearman’s rho = 0.52, p < 0.0001, **Figure 4**), and for motifs predicted to cause a ‘strong’ stall (Spearman’s rho = 0.54, p < 0.0001, **Figure S3**). Proteins whose median CUB rank was in the top percentile harbored an order of magnitude fewer di-prolyl motifs per 100 amino acids (AA) than proteins within the bottom CUB percentile (0.002 vs. 0.014 motifs per 100 AA, GC-controlled). These findings place similar observations from *E. coli* [29] into a broad phylogenetic context.

**Figure 4:**
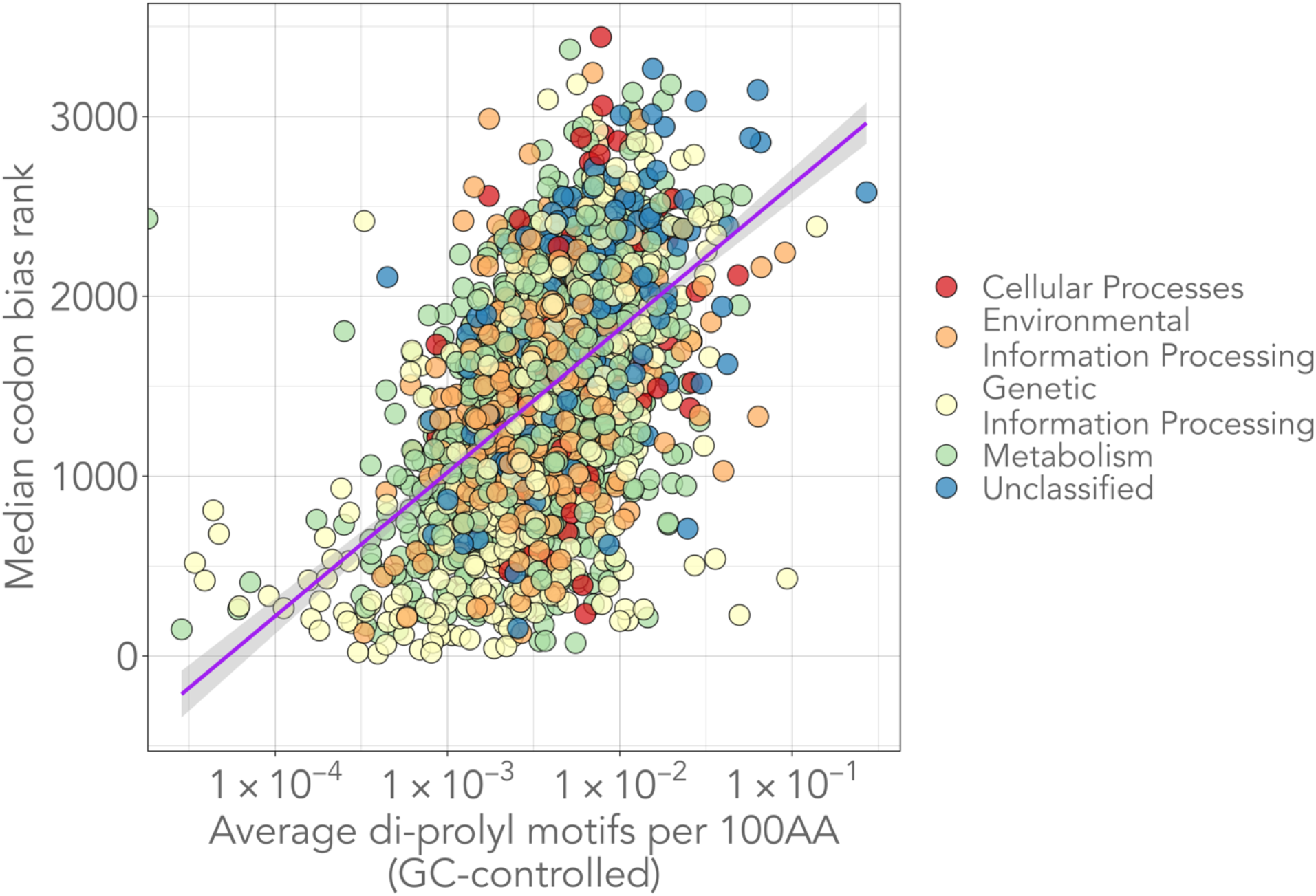
Highly expressed proteins contain fewer di-prolyl motifs. Proteins expected to be highly expressed (based on CUB) contain fewer di-prolyl motifs when protein length and gene GC content are controlled for (Spearman’s rho = 0.52, p < 0.0001). Each circle represents the average incidence of di-prolyl motifs within one protein (KEGG KO) across all genomes it was identified in. The colors represent the coarse-level function of the KEGG Orthology (KO) group to which each protein belongs. We only included common proteins (present in at least 25% of genomes) in this analysis to reduce bias towards rare proteins. The purple line is a linear regression line, and the shaded area represents the 95% confidence area. The horizontal axis is plotted on a logarithmic scale.

While highly expressed proteins were generally depleted in di-prolyl motifs, we wondered whether this association would be especially pronounced in fast-growing species. While we previously showed that fast-growing bacteria encode fewer di-prolyl motifs than slow-growing bacteria in general (**Figure 3)**, this analysis did not distinguish proteins based on their expected expression level. We hypothesized that fast-growing species would have fewer di-prolyl motifs in proteins that are likely to be highly expressed, because the ribosomal stalling that these motifs can cause is more severe in proteins with high translation initiation rates [16], and because selective pressure on translational speed increases with growth rate.

To validate this hypothesis, we identified for each genome those genes expected to be highly expressed, i.e., genes whose CUB lies within the top one percent of all genes. We then calculated the average amount of di-prolyl motifs in the proteins these genes encode. While fast-growing species are expected to show high CUB primarily in genes related to growth and translation (ribosomal proteins, translation factors, etc), slow-growing species also show evidence of CUB in highly expressed genes, although perhaps less dramatically and to a greater extent in non-translation-related genes [60]. For example, in slow-growing cyanobacteria with doubling times between 6-18 hours CUB is especially high in genes related to photosynthesis [60]. Indeed, in slow growing species (predicted doubling times longer than five hours), only 19% of genes in the top CUB percentile are translation-related, as opposed to 34% in fast growing species (predicted doubling times shorter than five hours; **Figure S4**). Regardless, we found that on average, fewer di-prolyl motifs were encoded by genes predicted to be highly expressed in fast-growing species than in slow-growing species, independent of GC content and growth temperature (**PGLS G**, predicted doubling time p < 0.0001, b = 0.037).

### Di-prolyl motifs are enriched in disordered domains of serine-threonine kinases and other signaling proteins

Previous analyses revealed an association between bacteria with high cellular complexity and elevated levels of di-prolyl motifs. Bacteria capable of multicellular behavior encoded more di-prolyl motifs independent of growth traits and growth temperature (**PGLS D**, p ≤ 0.01, b_unicellular_ = −0.015), with genomes from the Myxococcota containing by far the most di-prolyl motifs in our dataset (**Figure 1**). Proline-rich motifs are common in the binding domains of signaling proteins [13], a class of proteins crucial to orchestrating complex, multicellular lifecycles [61]. Because of these associations, we wondered whether multicellular bacteria contain more di-prolyl motifs purely because they encode more signaling proteins that help maintain and regulate multicellular organization.

To answer this question, we re-calculated the di-prolyl motifs encoded by the genomes in our dataset, excluding motifs present in proteins categorized by KEGG as either ‘Signal Transduction’ or ‘Protein Kinases’. We then analyzed these revised motif counts with a PGLS model and found multicellular bacteria still encoded significantly more di-prolyl motifs than unicellular bacteria (**PGLS H**, p ≤ 0.01, b_unicellular_ = −0.015). We note, however, that our observations are likely affected by the limitations of KEGG annotations and annotations for bacterial genomes in general [62]. Specifically, many genes in our dataset are not annotated in KEGG (49.4% on average), and multicellular genomes harbor significantly more unannotated genes than unicellular genomes (62.6% on average are unannotated by KEGG; phylogenetic ANOVA p ≤ 0.001). Thus, it is likely that our analysis does not actually encompass all signaling proteins, especially for genomes of poorly studied species [62].

While we found that di-prolyl motifs located in KEGG-annotated signaling proteins were not responsible for the elevated numbers of these motifs in multicellular bacteria, signaling proteins did contain significantly more di-prolyl motifs in multicellular bacteria (phylogenetic ANOVA, p ≤ 0.001). A large proportion of these motifs were in protein kinases, and specifically in serine-threonine kinases. Serine-threonine kinases are involved in signal transduction pathways in eukaryotes and some bacteria, and work by phosphorylating specific sites on target proteins [61, 63]. Such signaling can involve extensive cross-phosphorylation between kinases, modulating their downstream activity in signaling cascades [63]. Serine-threonine kinases regulate developmental processes in the Myxococcota [64, 65], and their prevalence within a genome is associated with increasing cellular complexity [61]. Multicellular bacteria contain significantly more serine-threonine kinases than unicellular bacteria (**Figure S5**, phylogenetic ANOVA p ≤ 0.001), and these kinases contain many di-prolyl motifs, especially within the Myxococcota. For example, *Stigmatella erecta* contains 530 di-prolyl motifs spread across the 67 serine-threonine kinases encoded within its genome. Di-prolyl motifs in this species are enriched 45-fold in serine-threonine kinases, such that 3.66% of di-prolyl motifs occur in a set of proteins that make up just 0.08% of its proteome.

The phosphorylation sites of protein kinases are predominately located within intrinsically disordered regions (regions without stable three-dimensional structure—IDRs), where phosphorylation can trigger transitions between disorder and order [66]. Notably, proline rich sites with high propensity towards forming polyproline II helices (PPII) are evolutionarily conserved at intrinsically disordered phosphorylation sites, where phosphorylation may ‘tune’ a protein’s propensity to adopt PPII structure [12]. These connections between proline rich PPII sites, phosphorylation, and IDRs led us to wonder whether the di-prolyl motifs within serine-threonine kinases were preferentially located within IDRs of these proteins, and thus potentially involved in transitions related to phosphorylation.

To answer this question, we used the disorder prediction software IUPred2A [45] to identify disordered regions in all serine-threonine kinases within our dataset. While on average only 27% of residues within the 6552 serine-threonine kinases in our dataset were disordered, 72% of di-prolyl motifs were located within these disordered regions, a significant enrichment (χ^2^ test p < 0.0001, **Figure S6**). In one extreme case, a single serine-threonine kinase from the myxobacterium *Stigmatella erecta* contained 41 di-prolyl motifs, of which 95% were located within disordered regions (**Figure 5**). While multicellular species contained more serine-threonine kinases than unicellular (**Figure S5**), and more di-prolyl motifs within these proteins (phylogenetic ANOVA p < 0.05; multicellular serine-threonine kinases contain 5.1 di-prolyl motifs on average vs 2.8 motifs in unicellular kinases), the di-prolyl motifs of both unicellular and multicellular serine-threonine kinases were equally enriched among disordered regions (71% of multicellular motifs occur in IDRs vs 72% of unicellular motifs). Expanding on these findings, we identified four additional common signaling proteins (encoded by 25% of genomes in our dataset) whose di-prolyl motifs were significantly enriched within disordered regions (χ^2^ test Bonferroni corrected p < 0.001, **Figure S6**). These findings indicate that the enrichment of di-prolyl motifs within IDRs may be a general feature of kinases and signaling proteins.

**Figure 5:**
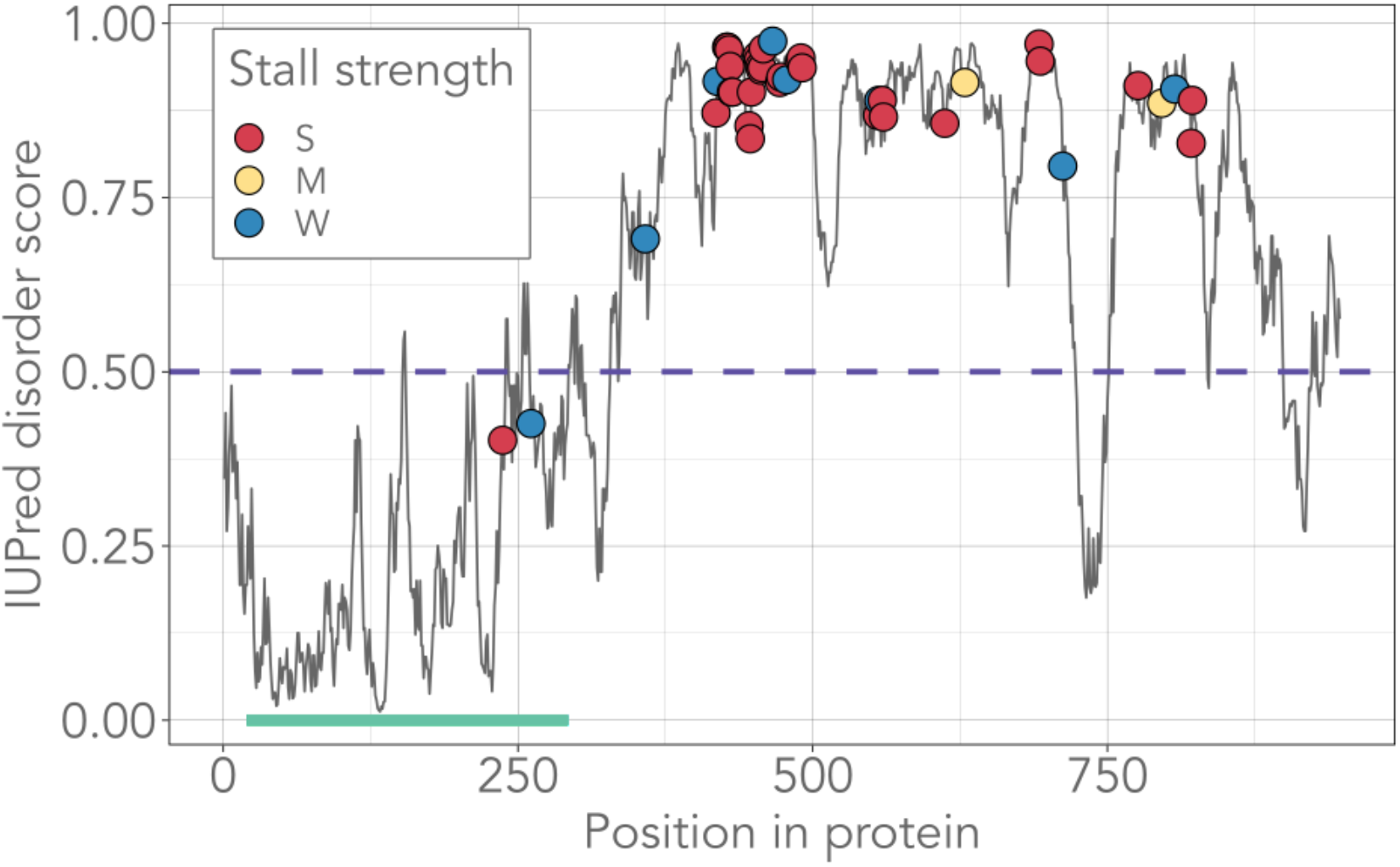
95% of di-prolyl motifs (39/41) within a single serine-threonine kinase from the myxobacterium *Stigmatella erecta* fall within intrinsically disordered regions. Each circle represents a single di-prolyl motif within the protein, with color indicating the predicted severity of the resulting stall, from strong (S) to medium (M) to weak (W). A region is predicted to be intrinsically disordered if the IUPred2A disorder score is greater than or equal to 0.5, as indicated by the dashed purple line. The green rectangle at the bottom of the figure indicates the PFAM protein kinase domain (PF00069) which is congruent with the set of ordered residues in this protein, as expected. The protein is encoded by IMG gene 2695004422 in IMG taxon oid 2693429888. The figure design is inspired by default IUPred2A plots (Erdos and Dosztányi, 2020).

## Discussion

Research involving EFP and di-prolyl motifs has largely focused on individual species [24, 25, 27, 29] or proteins [67]. In contrast, we surveyed these motifs in a broad range of genotypically and phenotypically diverse bacteria. While the exact effect of di-prolyl motifs on translational rate has not been experimentally tested in every species we studied, di-prolyl motifs have been shown to cause ribosomal stalling in all three domains of life, consistent with the near universal distribution of EFP and EFP homologs [17]. Likewise, every genome in our dataset encoded EFP, with one exception (see Methods). Because proline codons are cytosine rich (CCU, CCC, CCA, and CCG), it is not surprising that the occurrence of di-prolyl motifs is strongly associated with a genome’s GC content (**Figure 1**). We thus focused on patterns of di-prolyl motif occurrence that cannot be explained by GC content alone. We found such patterns in three groups of bacteria: slow-growing species, thermophilic species, and multicellular species.

Ribosomal stalling caused by di-prolyl motifs is resolved by bacterial elongation factor P (EFP). Without EFP, translational speed is reduced and proteins containing di-prolyl motifs can be under-expressed, leading to adverse consequences [15]. While EFP reduces proline-induced ribosomal stalling and its impact on translation speed, studies measuring ribosomal pausing show that EFP cannot completely resolve these stalls—even in the presence of EFP ribosomes still pause at proline residues [15, 28]. Such stalling can be especially detrimental in fast-growing species, which can be sensitive to minute disruptions in translational rate [2]. For example, in *E. coli*, even a single sub-optimal codon in a lowly expressed gene can impact fitness [68]. *E. coli* also appears to be sensitive to the presence of di-prolyl motifs, because its genome encodes fewer di-prolyl motifs than expected by chance, and fewer still in genes encoding highly expressed proteins [29].

We wanted to find out whether these species-specific observations translate to a broad range of bacteria. We were especially interested in whether the prevalence of di-prolyl motifs would be associated with species-specific growth rates. Indeed, we found that fast-growing species encode significantly fewer di-prolyl motifs than slow-growing species (**PGLS D**, **Figure 3**). This relationship holds whether we estimate growth rate indirectly from codon usage bias (**PGLS D, Figure 3**), or directly from experimental measurements (**PGLS F, Figure S2**). Additionally, while proteins expected to be highly expressed generally encode fewer di-prolyl motifs (**Figure 4 and S3**), this trend is exacerbated in fast-growing species (**PGLS G**). These findings are based on analyses of variance that correct for phylogenetic relatedness and allow the impact of co-correlated traits to be quantified independently (PGLS, **Supplemental Table 1**).

Several highly expressed proteins critical to cell function contain di-prolyl motifs that cannot be removed without a resulting loss of function. For example, the valine tRNA synthetase ValS contains a poly-proline motif that is highly conserved, critical to charge tRNA^Val^ efficiently, and important to prevent its mischarging with threonine [67]. This implies that some di-prolyl motifs are maintained in the face of negative selection pressure because the benefit of their specific biochemical activity outweighs any impact they may have on translational rate. If the genomes of some species can encode more di-prolyl motifs because of weaker selection on translational rate, these motifs may also acquire new and useful roles unrelated to their effect on protein translation.

One candidate for such a role is to stabilize proteins in the high temperature environments experienced by thermophilic bacteria. Proline residues can increase the thermostability of proteins by at least two mechanisms. First, their rigid structure reduces the degrees of freedom available to a protein [53]. Second, increasing the proportion of prolines and other hydrophobic residues enhances thermostability by reducing accessibility to the protein core [53]. Thermophiles have smaller genomes and shorter proteins than mesophiles [54]. However, when we controlled for the proportion of their genome that encodes proteins, we found that thermophiles encoded more di-prolyl motifs than mesophiles (**PGLS A**, **Figure 2A**). Thermophiles are thought to experience relaxed selection on translation rate because catalysis naturally proceeds more rapidly at higher temperatures [5]. In other words, thermophiles generally grow much more quickly than their overall investment in maximizing translational rate would suggest. Relaxed selection on translational speed in thermophiles may allow prolines to exist where they would otherwise be detrimental for translational rate, and thus help stabilize a thermophilic proteome.

Another potential role for di-prolyl motifs exists in bacteria with complex, multicellular lifecycles, most notably the Myxococcota (**Figure 1**). We found that multicellular bacteria contained significantly more di-prolyl motifs than unicellular bacteria (**PGLS B, Figure 2B**). These organisms rely on signaling proteins to orchestrate their complex lifecycles [64, 65], which often contain proline-rich regions with their binding domains [12, 13]. While we found signaling proteins were not the sole source of the elevated levels of di-prolyl motifs within these species (**PGLS H**), these results are likely biased by the especially poor level of gene annotation in multicellular species (62.6% of genes in multicellular genomes are unannotated by KEGG vs. 49.6% for unicellular genomes).

Multicellular species do harbor disproportionately high numbers of the di-prolyl rich signaling proteins serine-threonine kinases compared to unicellular bacteria (**Figure S5**). These kinases contain an outsized proportion of di-prolyl motifs, which can be enriched up to 45-fold in some Myxococcota compared to their background occurrence within the proteome. Serine-threonine kinases play central roles in signaling between cells by modulating the activity of their target proteins through phosphorylation [63] and have been linked to cellular complexity in multicellular bacteria [61].

Bacterial kinases are known to cross and auto-phosphorylate, creating extensive signaling networks [63]. The phosphorylation sites of kinases are enriched in intrinsically disordered regions (IDRs), where phosphorylation can trigger changes in three-dimensional structure that alter downstream activity [66]. Interestingly, proline rich regions with high propensity towards forming polyproline II helices (PPII) also occur primarily within IDRs [69], and are evolutionarily conserved at phosphorylation sites [12]. Following these connections, we found that di-prolyl motifs within serine-threonine kinases are enriched among IDRs, as illustrated by a specific example in **Figure 5** and more generally in **Figure S6**. Furthermore, we found that the di-prolyl motifs of four other common signaling proteins (present in > 25% of our genomes) were significantly enriched among IDRs (**Figure S6**). Three of these signaling proteins are known to be phosphorylated (response regulator RegA [70], sensor kinase CheA [71], and an OmpR family sensor kinase [72]). Phosphorylation can have a dramatic effect on local bias towards PPII structures, in effect ‘tuning’ a protein’s propensity towards adopting a PPII structure [12, 73]. As PPII structures commonly form the binding domains of signaling proteins [12, 13], a connection between phosphorylation, PPII formation (or collapse), and the modulation of signaling protein activity is appealing. However, verifying these connections and determining their biological significance remains a task for future work.

Our observations are consistent with previously drawn connections between cellular complexity and polyproline motifs, with more complex organisms containing higher numbers of such motifs [10]. This connection may also be related to the increasing number of IDRs in complex genomes, which in turn are linked to the importance of post-translational modification for complex signaling pathways [69, 74]. The multicellular bacteria in our dataset are generally slow-growing, with an average predicted doubling time of 8.4 hours, and the Myxococcota are especially slow-growing, with an average predicted doubling time of 12.5 hours. As a result, selection on translational rate is weaker in these species, which may allow prolines to accumulate where they would otherwise be discouraged. Indeed, it could be informative to interrogate possible regulatory functions of EFP in these species, as an important class of signaling proteins in multicellular bacteria is enriched in di-prolyl motifs (**Figure 5**), and EFP may influence the expression of these proteins.

In conclusion, our observations suggest active selection against di-prolyl motifs in a broad range of fast-growing species, and in highly expressed proteins of such species. Such selection is likely driven by high selection pressure on optimizing translational rate. Wherever such selection is relaxed, di-prolyl motifs may be free to proliferate and take up new roles. One of these roles may be to ensure proteome stability in thermophiles. Another may be to help cells in simple multicellular prokaryotes communicate. However, the causal role of di-prolyl motifs in any of these processes is unclear. For example, did multicellular bacteria emerge from slow-growing unicellular bacteria, where a high incidence of di-prolyl motifs facilitated kinase-based cell signaling and helped establish multicellularity? Or did multi-cellular bacteria emerge from fast-growing unicellular bacteria, such that their reduced growth rate, kinase-based signaling, and the importance of di-prolyl motifs emerged only secondarily? These and other questions about the biological functions and evolutionary origins of di-prolyl motifs provide exciting directions for future work.

## Data availability

All genomes used in this study are publicly available and were downloaded from JGI’s IMG database [35]. Taxon IDs corresponding to every genome included in our dataset are listed in **Supplemental Dataset 1**, along with the genomic characteristics calculated for this study. Results from analyses of individual proteins are presented in **Supplemental Dataset 2** and results of all PGLS models are listed in **Supplemental Table 1**. R scripts and all files needed to reproduce these analyses are available at https://github.com/tessbrewer/proline_project.

## Supporting information

Supplemental Figures + File Descriptions

## Acknowledgements

We acknowledge funding from the European Research Council under Grant Agreement No. 739874, as well as from Swiss National Science Foundation grant 31003A_172887, and from the University Priority Research Program in Evolutionary Biology at the University of Zurich. We thank members of the Wagner lab for helpful discussions, particularly Andrei Papkou and Pouria Dasmeh, and Michael Engel for figure design input.

## Author contributions

TEB and AW conceived and designed the project. TEB performed all computational analyses. TEB and AW wrote the paper.

