## Supplemental Figures + File Descriptions for "Translation stalling proline motifs are enriched in slow-growing, thermophilic, and multicellular bacteria"

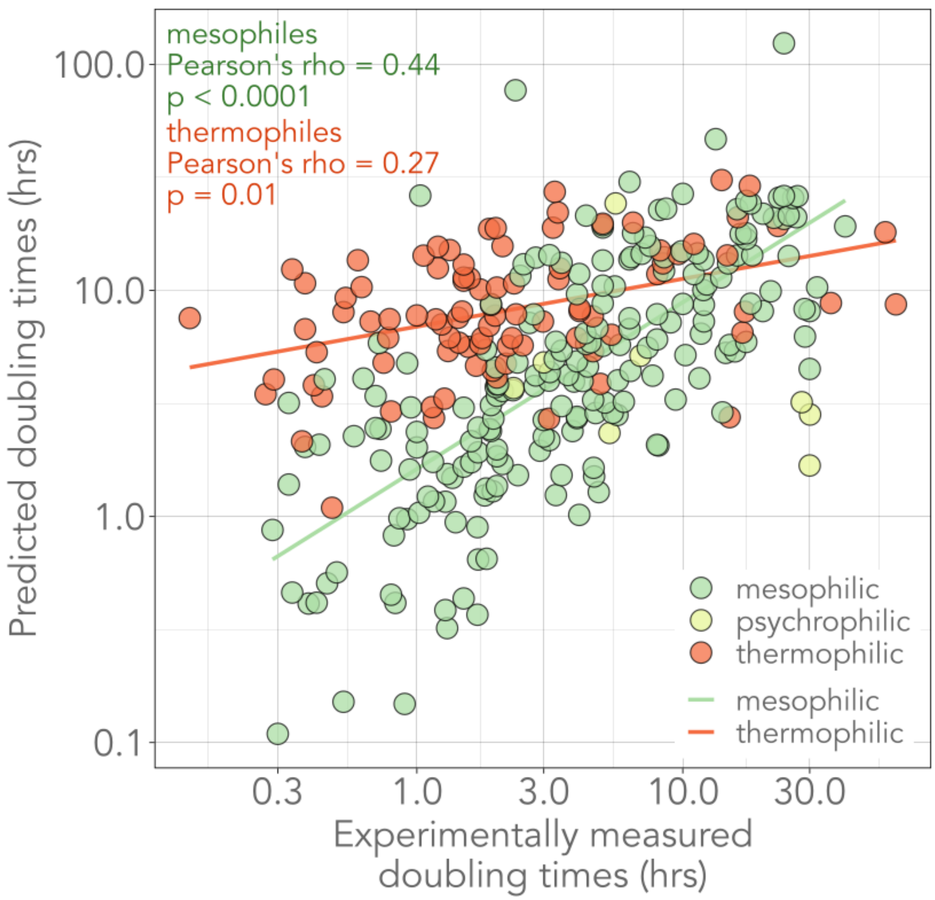


**Figure S1: CUB-based estimated doubling times correlate well with experimentally measured doubling times, especially in mesophiles.** Doubling times estimated using the CUB based R package gRodon (Weissman *et al.*, 2021) are moderately correlated with experimentally measured doubling times (Pearson's rho = 0.33, $p < 0.0001$, $n = 301$). This correlation improves when only mesophilic species are considered (Pearson's rho = 0.44, $p < 0.0001$, $n = 202$). Measured doubling times are known to be misestimated by CUB-based methods in thermophiles and psychrophiles (Vieira-Silva and Rocha, 2009; Weissman *et al.*, 2021). The horizontal and vertical axes are plotted on a logarithmic scale.

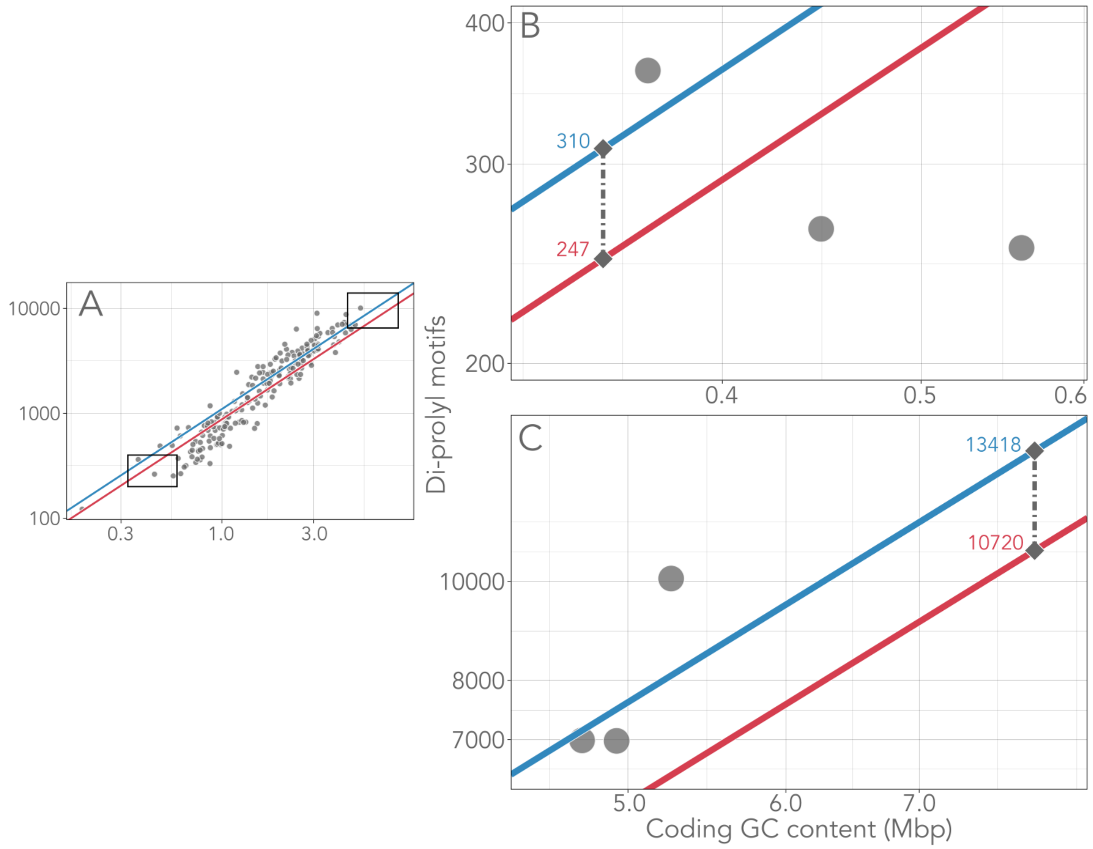


**Figure S2: Slow-growing bacteria still encode more di-prolyl motifs than fast-growing bacteria when experimentally measured doubling times are used in place of CUB-predicted doubling times. A:** We identified experimentally measured doubling times for 202 mesophilic species (6.2% of our full dataset) and used these values in a PGLS model in-place of predicted doubling times. The results are consistent with the PGLS analysis of our full dataset (**Figure 3**). Although GC content is the primary determinant of the number of di-prolyl motifs in a genome (**PGLS F** $p < 0.0001$, $b = 1.204$; b = regression slope), measured doubling time and tRNA gene copy number also have a significant impact (**PGLS F**; measured doubling time $p < 0.001$ $b = 0.034$, tRNA gene copies $p < 0.05$ $b = -0.108$). rRNA gene copy number is insignificant in this smaller dataset ($p = 0.315$). As in Figure 3, the two diagonal lines represent the number of PGLS-predicted di-prolyl motifs, as calculated from the growth-associated traits of two representative organisms in our dataset. Specifically, the upper blue line represents a prediction based on one of the slowest growing species (*Methylomagnum ishizawai*; experimentally measured doubling time = 24 hrs, tRNA gene copies = 51) while the lower red line represents a prediction based on one of the fastest growing species (*Propionigenium maris*; experimentally measured doubling time = 0.3 hrs, tRNA gene copies = 103). In this reduced dataset, the slow-growth related traits of *Methylomagnum ishizawai* result in a 25% predicted increase in di-prolyl motifs. At low GC content (**B: enlarged view of the lower left box in A**), the total impact of growth-associated traits is low (calculated net increase of only 63 di-prolyl motifs at 0.35 Mbp GC content), while at high GC content (**C: enlarged view of the upper right box in A**), the total net increase is substantial (calculated net increase of 2698 di-prolyl motifs at 8 Mbp GC content). The horizontal and vertical axes are plotted on a logarithmic scale.

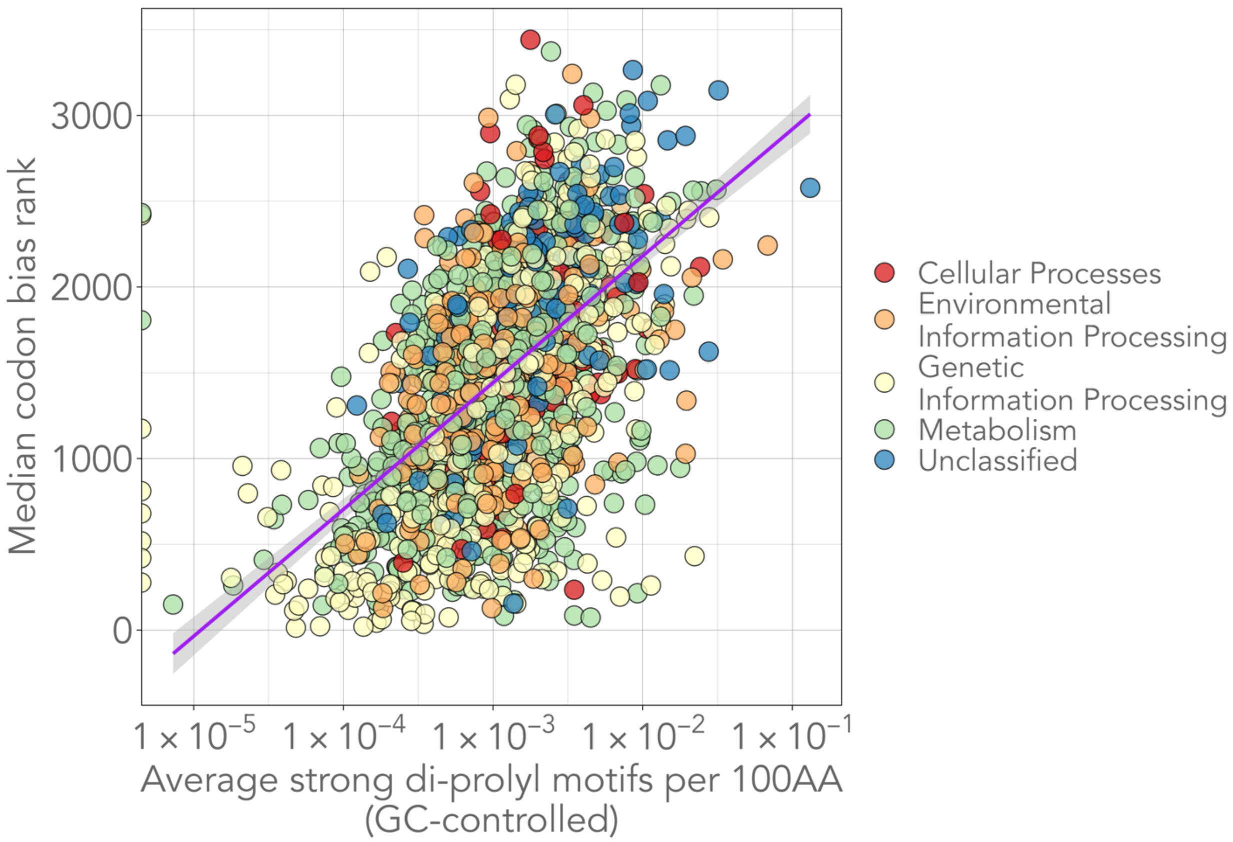


**Figure S3: Proteins optimized for translational efficiency contain fewer 'strong' di-prolyl motifs**. The number of 'strong' di-prolyl motifs in a protein decreases as the protein's predicted expression level increases, when protein length and gene GC content are controlled for (Spearman's rho = 0.54, $p < 0.0001$). Each circle represents the average incidence of strong di-prolyl motifs within one protein across all genomes it was identified in. The colors represent the coarse-level function of the KEGG Orthology (KO) group to which the protein belongs. We only included common proteins (present in at least 25% of the genomes in our dataset) in this analysis to reduce any bias towards rare proteins. The purple line is a linear regression line, and the shaded area represents the 95% confidence area. The horizontal axis is plotted on a logarithmic scale.

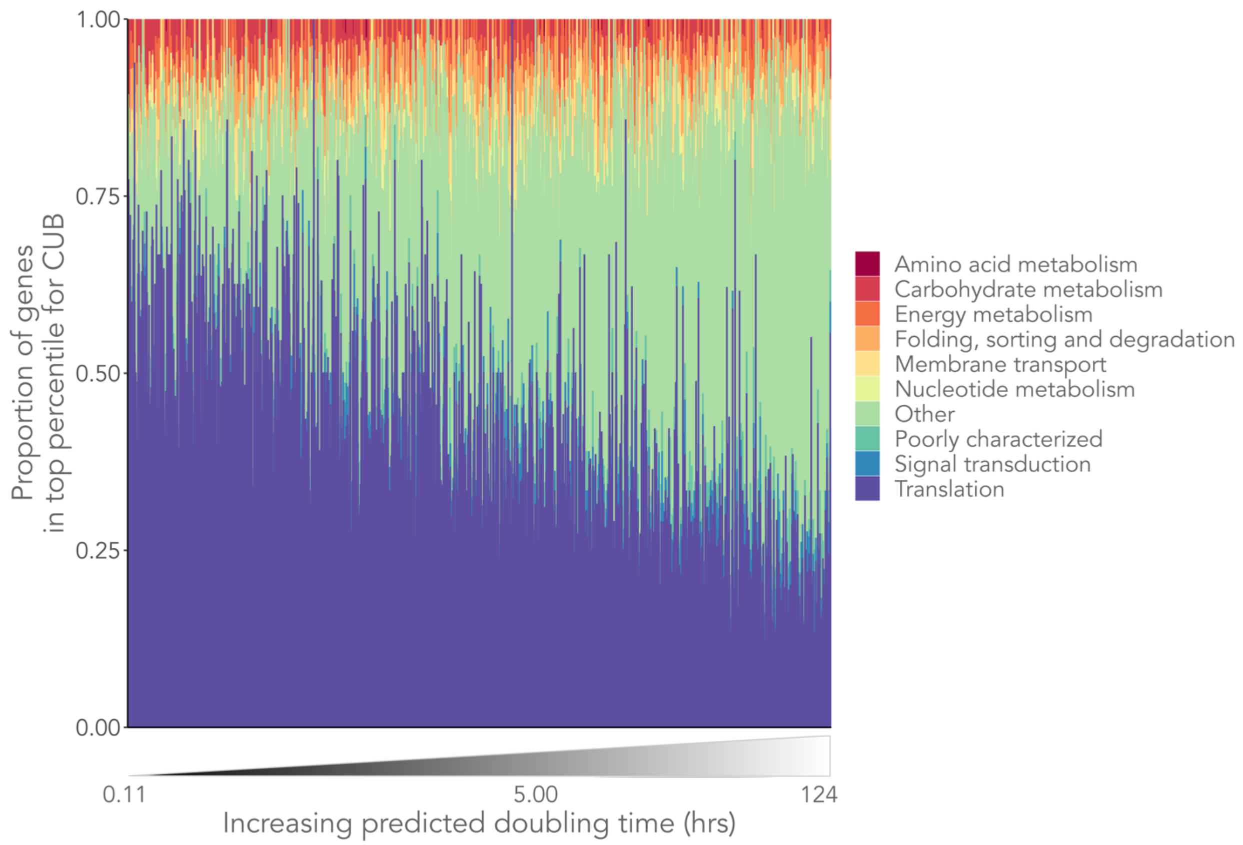


**Figure S4: Slow-growing bacteria have fewer translation-related genes within their most CUB optimized genes.** On average, 34% of genes optimized for high expression (within the top percentile for CUB) are translation related in fast-growing species (predicted doubling time < 5 hours) vs. 19% for slow growing (predicted doubling time > 5 hours) species. Each bar represents genes expected to be highly expressed within a single species. The colors show the distribution of these genes across B-level KEGG categories. Only the 9 most abundant B-level KEGG categories are shown for clarity. Genes belonging to other categories or those without a KEGG annotation are grouped together as ‘Other’.

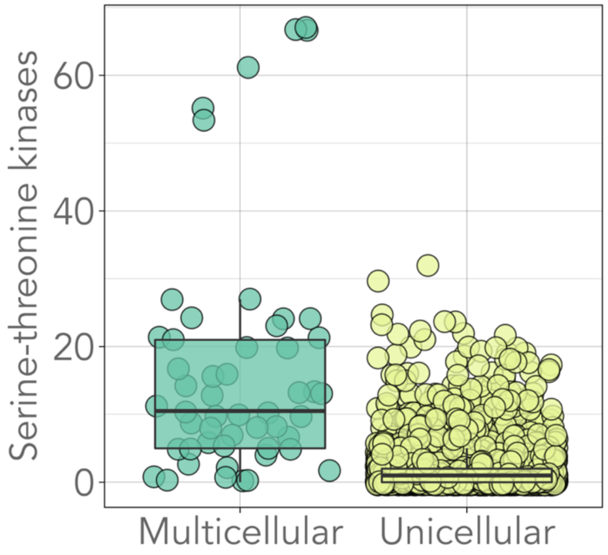


**Figure S5: Multicellular bacteria encode more serine-threonine kinases than unicellular bacteria**. (Phylogenetic ANOVA $p \leq 0.001$). The box spans the first and third quartiles, while the horizontal line within represents the median. Each circle represents the number of serine-threonine kinases (specifically from KEGG Orthology group K08884) within one genome. All six multicellular genomes encoding more than 50 serine-threonine kinases hail from the phylum Myxococcota.

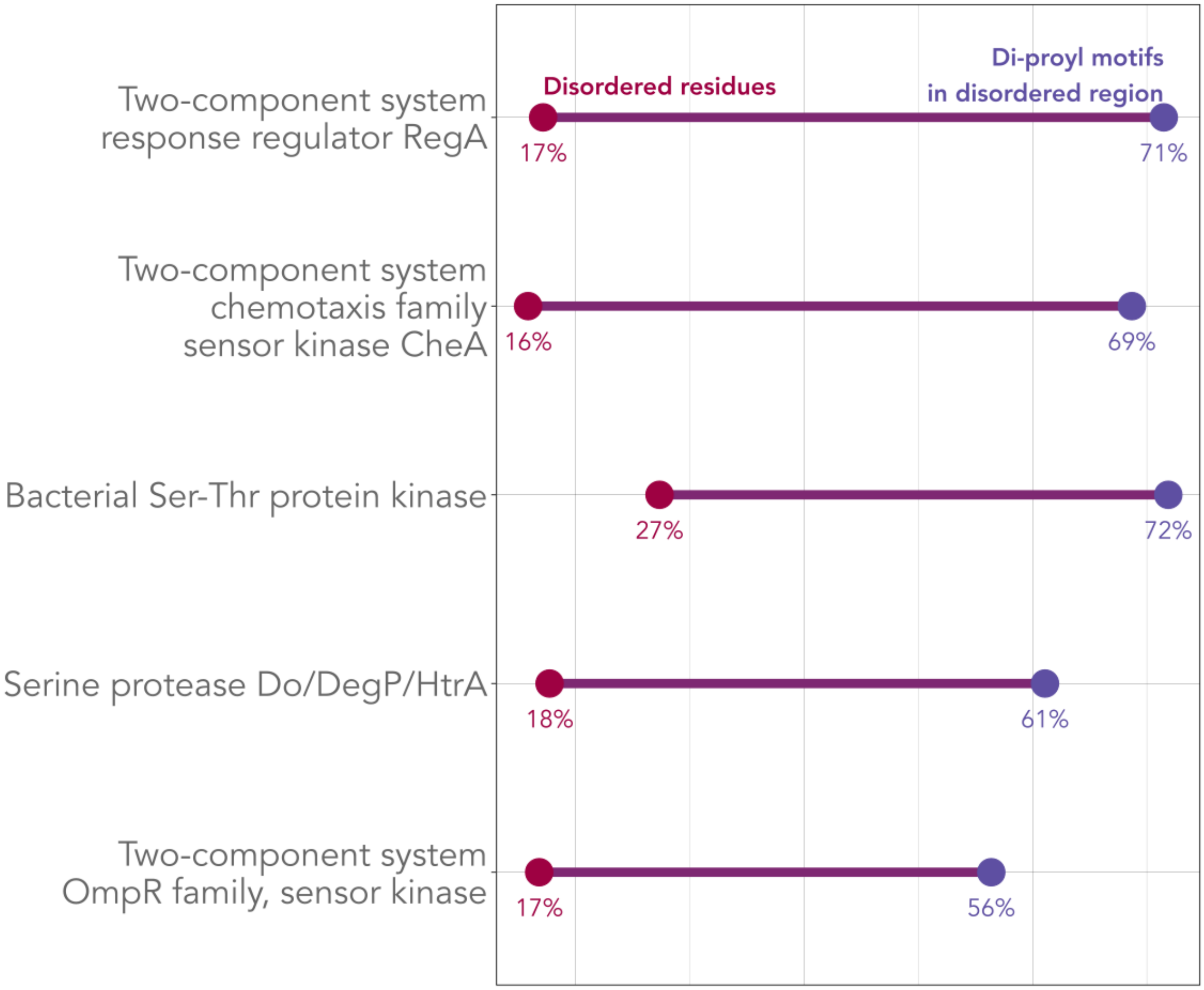


**Figure S6: Di-prolyl motifs are significantly enriched in intrinsically disordered regions of five common signaling proteins.** The figure shows data from five common signaling proteins (present in > 25% of genomes in our dataset) which contain di-prolyl motifs that are significantly enriched in intrinsically disordered regions (IDRs) ($\chi^2$ test Bonferroni corrected $p < 0.001$). Red circles indicate the average percent of amino acid residues that are predicted to be disordered within these proteins (IUPred2A disorder score $\geq 0.5$), while the blue circles indicate the percentage of di-prolyl motifs that occur within these disordered regions (IUPred2A disorder score $\geq 0.5$, averaged over the five amino acids making up the motif). Four of these proteins are known to be phosphorylated (RegA, CheA, Ser/Thr-protein kinases, OmpR family kinases). Only proteins that frequently contain di-prolyl motifs (di-prolyl motif is present in > 50% of genomes containing the protein) and with at least 30% enrichment in di-prolyl motifs located in IDRs are shown for clarity. The KEGG Orthology classifications for all proteins in descending order are: K15012, K03407, K08884, K04771, K02484.

**Supplemental Table 1:** Details of all models presented in paper.

**Supplemental Dataset S1:** Dataset including information on all genomes used in these analyses.

**Supplemental Dataset S2:** Dataset including information on all proteins used in these analyses. Based on KEGG orthology.
